# Primary productivity and N_2_-fixation in the eastern Indian Ocean: bottom-up support for an ecologically and economically important ecosystem

**DOI:** 10.1101/2025.09.25.678645

**Authors:** Sven A. Kranz, Jared M. Rose, Michael R. Stukel, Karen E. Selph, Natalia Yingling, Michael R. Landry

## Abstract

Oligotrophic regions of the global ocean are characterized by strong nutrient limitation, low standing phytoplankton biomass, and highly efficient nutrient recycling. We quantified nutrient inventories, primary productivity and N_2_ fixation during the BLOOFINZ-IO expedition (February 2022) in the Argo Basin located in the eastern Indian Ocean, the sole known spawning ground for Southern Bluefin Tuna. Surface nitrate concentrations were near depletion (<0.02 µmol L^-1^), with low but persistent residual phosphate (P) concentrations suggesting nitrogen as the major limiting nutrient. Depth-integrated net primary production (NPP), from ^14^C-based in-situ incubations during 4 Lagrangian cycles, averaged ∼460 mg C m^-2^d^-1^, in good agreement with satellite-based NPP estimates. Nitrogen fixation provided a consistent new nitrogen source, contributing ∼16% to local NPP in the upper euphotic zone.

Gross primary production (GPP), derived from fast-repetition-rate-fluorometry-based electron transport estimates, revealed significant autotrophic respiration losses, with GPP:NPP ratios averaging ∼1.8, consistent with metabolic costs under nutrient limitation. Net community production (NCP), estimated from O_2_/Ar ratios, remained positive across all cycles, averaging ∼20% of NPP in the upper 30 m. This result, in combination with N_2_ fixation measurement indicates that N_2_ fixation supports most of the export production in this region. Together, the multi-method approach revealed a recycling-dominated ecosystem affected by episodic mixing events, where primary productivity is maintained primarily through efficient nitrogen recycling and physiological photoacclimation. These results provide a comprehensive baseline of bottom-up support on ecosystem productivity for the Argo Basin for assessing future climate-driven changes in stratification, nutrient cycling, and food-web dynamics.

## 1. Introduction

The eastern Indian Ocean, located between northwestern Australia and directly downstream of the Indonesian Throughflow, is a region of global significance for regulating heat transfer from the Pacific to Indian Ocean (Lee et al., 2015; Desbruyeres et al., 2017) and is the only known spawning region for Southern Bluefin Tuna (SBT) (Farley and Davis, 1998). However, it remains a sparsely studied area regarding its ecological and biogeochemical processes. The BLOOFINZ (Bluefin Larvae in Oligotrophic Ocean Foodwebs, Investigations of Nutrients to Zooplankton) project investigated the region during the peak SBT spawning season in January-February 2022 to assess its potential vulnerabilities to future climate change. Quantification of gross-, net- and net community-productivity and its support by nitrogen (N) sources are central to understanding the energy available for higher trophic levels. In the present study, we focus on primary production, N_2_ fixation and associated photophysiologial measurements conducted during four multi-day Lagrangian experiments.

Despite low rates of production per unit area, oligotrophic systems contribute an estimated 30–40% of total global marine primary production (Behrenfeld and Falkowski, 1997; Field et al., 1998). Primary productivity depends on efficient recycling of N through ammonification (Dugdale and Goering, 1967). In addition, N_2_ fixation can play a significant role in supporting new production (Raes et al., 2015; Tang et al., 2019). Due to the need to retain recycled nutrients for continuing production, only a small fraction of primary production is usually exported out of the euphotic zone (Buesseler et al., 1998; Henson et al., 2019). Deep chlorophyll maxima (DCM) also hide a substantial portion of the phytoplankton community and their dynamics from satellite remote sensing, raising questions of how general algorithms relate to depth-resolved measurements of net primary production (NPP) from ^14^C uptake or to high-resolution quantification of gross primary (GPP) and net community productivity (NCP) (Robinson et al., 2009; Quay et al., 2010; Hamme et al., 2012; Schuback et al., 2015; Teeter et al., 2018; Kranz et al., 2020).

In this study, we applied a multi-method approach to quantify nutrients, productivity metrics and N_2_ fixation in SBT spawning waters overlying the 5000-m deep Argo Abyssal Plain (hereafter, Argo Basin) off NW Australia (Fig.1). We directly compare NPP assessments from *in situ* incubations and satellite products for four multi-day Lagrangian experiments. From continuous surface measurements of O_2_:Ar, Fast Repetition Rate Fluorometry (FRRf) and photophysiological parameters in transect sampling across the basin, we derive carbon-based estimates of NCP, GPP and production relationships. From direct experimental measurements of N_2_ fixation rates, we establish that diazotrophy is the main contributor to the new N required to balance atypically high export fluxes. Our overall goals were to evaluate the magnitude and variability of autotrophic production and N supply to this key region and to establish baseline metrics for understanding future responses of the system to climate change.

## 2 Material and Methods

### 2.1 General sampling plan

The Argo Basin was sampled on BLOOFINZ cruise RR2201 of *R/V Roger Revelle* between 31 January and 3 March 2022 (Fig. 1). We first conducted an east-to-west transect across the central basin, then sampled south to begin the first of four Lagrangian experiments (hereafter, “cycles”) in the southern basin. Following the cycle experiments, two additional transects were sampled before departing the basin, first east-to-west at 15°S, then west-to-east at 13.5°S. During each cycle, sampling and incubations were done on a repeated daily schedule for 3-5 days following a satellite-tracked drifter with a 3-m drogue centered at 15-m depth (Landry et al., 2009; Stukel et al., 2015). Seawater samples for nutrients, phytoplankton and experimental incubations were collected with a 12-place CTD rosette of 10-L Niskin bottles. Incident Photosynthetically Active Radiation (PAR) was measured continuously by the ship’s meteorological system and a PAR sensor on CTD profiles. During both transect and cycle experiments, continuous measurements of mixed-layer O_2_:Ar ratios, fluorometric chlorophyll *a* (Chl*a*) and FRRf were taken by instruments plumbed into the ship’s seawater intake delivered at ∼4 L min^-1^ by a diaphragm pump (Graco Husky 1050e).

**Figure 1:**
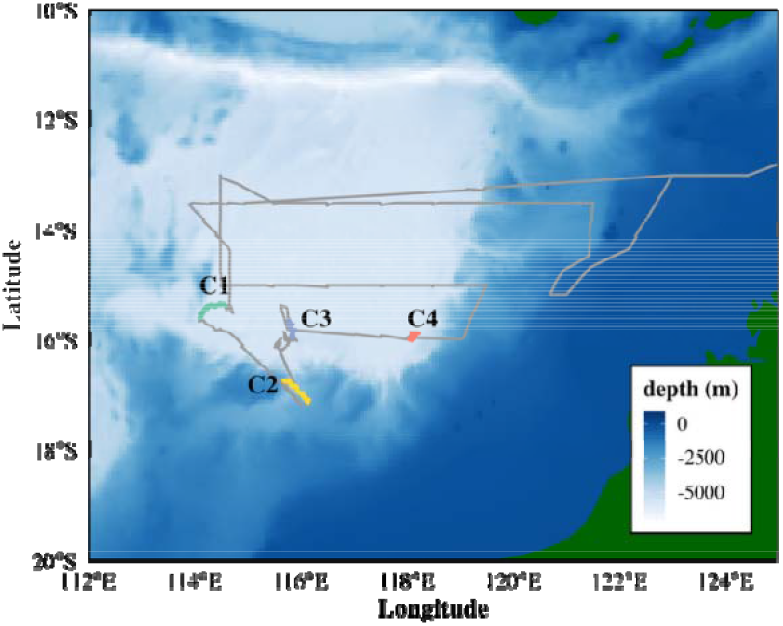
Map of the study area as a bathymetry with cruise track (in grey) and cycle location (in colors).

Water samples for extracted Chl*a* analyses were taken from CTD profiles, filtered onto GF/F filters, extracted in 90% acetone for 24 h and measured with a 10AU fluorometer (Turner Designs). Surface Chl*a* values were also continuously measured by a fluorometer. However, underway data between February 14 and 23 was unusable due to biofouling. Nutrient samples (50 mL) were filtered directly from CTD bottles through acid-cleaned 0.2-µm Acropak in-line capsules (Pall-Gelman) into 3X-rinsed Nalgene HDPE bottles and immediately frozen at -20 °C. Concentrations of nitrate, nitrite, ammonium, phosphate and silicate concentrations were measured with a Seal AutoAnalyzer 3 at the Scripps Institution of Oceanography Data Facility.

### 2.2 Net Primary Productivity (NPP) on in-situ array and satellite derived NPP

Net primary production was measured by ^14^C-bicarbonate uptake (NPP_14C_) in *in-situ* incubations following Morrow et al. (2018). For each cycle day, water was collected at six depths spanning the euphotic zone on the 02:00 (local) CTD cast and transferred gently via silicon tubing into acid-washed 280-mL polycarbonate incubation bottles that had been pre-soaked with Milli-Q water and rinsed three times with station water. For each depth, three replicate bottles and one dark bottle were spiked with 10 µCi H^14^CO ^-^ and incubated for 24 h (beginning at 04:00 local) in mesh bags attached at the collection depth. Following incubation, samples were processed under dim red light. After a subsample (100 µL) was taken for total radioactivity, bottle volumes were filtered onto GF/F filters, acidified with 0.5 mL of 10% biological-grade HCl, and degassed for 24 h. The filters were subsequently placed into scintillation cocktail and analyzed with a liquid scintillation counter. NPP_14C_ was calculated according to (Steemann Nielsen, 1952) as:

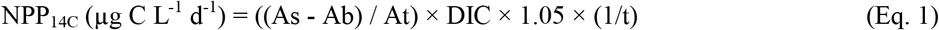

where As = sample activity; Ab = blank activity; At = total bottle activity; DIC = dissolved inorganic carbon (25.5 µmol C L^-1^); and t = incubation time (1 day). Depth integrations were done in Rstudio by trapezoidal interpolation to 30 m and to the deepest sample measurement.

NPP_Sat_ was also estimated for the Argo region from 8-day composite NPP satellite data products of the CAFÉ model from Oregon State University (Silsbe et al., 2016). Satellite based estimates of NPP for the region had been described by (Kehinde et al., 2023). Here, we re-analyzed the data along the cruise track, focusing specifically on the Lagrangian cycles, and averaging the data over 1-min intervals based on the ship’s location.

### 2.3. N_2_ fixation on in-situ array

N_2_ fixation rates were measured using the ^15^N_2_ gas tracer technique following the bubble dissolution method described by (Mohr et al., 2010). Because of the extra setup time required, samples for these incubations were taken from the 22:00 CTD and thus differed from 02:00 (local) productivity samples, though were taken in close proximity to the drift array and assumed to have a similar plankton composition. For each of 6 depths, unfiltered seawater was directly transferred into triplicate acid cleaned 4.6 L polycarbonate bottles and kept in the dark until spiked with ^15^N_2_-containing water. To improve dissolution of ^15^N_2_ into the spiked inoculum, 1.2 L of the same water was degassed (Mohr et al., 2010) using a Pfeiffer vacuum pump and rigorously stirred with Teflon-coated stir bar. The partially degassed water was transferred by peristaltic pump into a 1 L tedlar gas sampling bag (Restek) with residual dissolved O_2_ measured during transfer with an optode system (Firesting, Germany). On average, 35% of O_2_ was retained indicating a similar residual ^14^N_2_. The bags were filled with 1.2 L degassed seawater, spiked with 35 ml of ^15^N_2_ gas (98–99 atom %, Cambridge Isotope Laboratories), and transferred to a automatic bubble disturbing setup which moved the bubble continuously for ∼1.5 h. The ^15^N_2_ enrichment of this inoculum was calculated to be ∼50%. Around 03:00, an hour before array deployment, ∼350 ml of the inoculum was added to each of three 4.6-L bottles, and the bottles were filled to capacity with natural seawater for a final ^15^N_2_ enrichment of around 7.5%. These bottles were subsequently incubated for 24 h on the in-situ array.

Post incubation, aliquots from each bottle were transferred into 50-125 ml serum bottles using a peristaltic pump, and 5-12 µl of HgCl was added to kill biological activity. These bottles were sealed with Teflon-coated septums for gas-tight seals and stored in the dark at room temperature. Residual water from the incubations was filtered onto precombusted GF/F under dim light, and filters were frozen at -20°C before being dried in a desiccator and packed into tin cups for analysis at the Stable Isotope laboratory. The ^15^N_2_ spike concentration was measured on shipboard with a Pfeiffer QMS 220 MIMS system. A long needle inserted into the serum bottle was connected by a microvolume gear pump to the membrane system (Bay instruments), and the ^15^N:^14^N gas ratio was measured and found to average 5.7% (slightly less than theoretically assumed). N_2_ fixation was calculated according to (White et al., 2020). Depth integrations were done in Rstudio by trapezoidal interpolation to 30 m and to the deepest sample measurement at each cycle day (max. 90m).

### 2.4. Net community production (NCP)

NCP estimates from O_2_:Ar measurements reflect oxygen production by photoautotrophs, respiration by photo- and heterotrophs and corrections for physical gas exchange processes assuming steady-state processes over the residence time of O_2_ (Craig and Hayward, 1987). Continuous underway measurements of O_2_:Ar were obtained by equilibrium inlet mass spectrometry (EIMS, Pfeiffer QMS 220) as described by (Kranz et al., 2020). Biological supersaturation of oxygen (ΔO_2_:Ar) was determined by normalizing the measured O_2_:Ar ratio to values obtained during regular (∼1 h every 8 h) calibrations with atmosphere-equilibrated seawater. Data were smoothed by 5-min running averaging, and calibration times were exchanged with correlated extrapolations of the prior and post calibration data. NCP rates were calculated as:

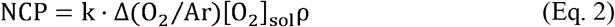

where k is the piston velocity for O_2_ gas exchange (k; m d^-1^) estimated from the square of satellite-measured wind speed and the temperature-dependent Schmidt number (Sc) for O_2_ (Wanninkhof, 2014). To incorporate the temporal dynamics of wind forcing and upper-ocean mixing, a time-weighted mean k was computed by incorporating the wind speed history and mixed-layer depth (MLD) (Reuer et al., 2007). [O_2_]_sol_ is the mixed-layer oxygen solubility, and ρ is the average ML density. Δ(*O*_2_ /Ar) is the biological oxygen signal defined by 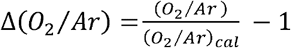.

NCP was calculated for two estimates of mixed layer depths (MLD_σθ_ and MLD_O2_) as well as the cycle average depth of 30m. Mixed-layer depth (MLD_σθ_) was determined from CTD profiles as the depth exceeding density at 5 m by 0.01 kg m^-3^. We also calculated MLD_O2_ as a 2% change in O_2_ concentration from a reference depth of 10 m (Castro-Morales and Kaiser, 2012). Both MLD estimates were smoothed by 3-d running means prior to the ventilation calculation to account for internal waves and other variations. NCP was calculated as mmol O_2_ m^-2^ d^-1^ and converted to NCP_C_ (mg C m^-2^ d^-1^) using a photosynthetic quotient of 1.3 and 12.01 g mol^-1^ for carbon.

### 2.5. Gross primary productivity and photophysiology using FRRf

Gross primary productivity (GPP) was estimated using Fast Repetition Rate fluorometry (FRRf) based on (Oxborough et al., 2012) and adapted for field applications following (Schuback et al., 2015; Schuback et al., 2016). Implementation of the method for this study closely followed (Kranz et al., 2020). Shipboard measurements were made from the ship’s seawater system using a bench-top FastAct 2+ Fast TRAKA instrument (Chelsea, UK). Photosynthesis versus irradiance (P vs. E) curves were run on a ∼30-min sampling interval using single-turnover flash sequences to resolve photosystem II (PSII) photophysiological parameters. For every 5-ml sample, water was first purged through the cuvette to remove old sample and acclimated to low light (12 µmol photons m^2^ s^-1^) for 10 min, then dark acclimated for 60 sec. Measured parameters included the dark/low light adapted maximum quantum yield of PSII (Fv/Fm), the functional absorption cross-section of PSII (σPSII), and the effective quantum yield under ambient light (Fq/Fm’). Additional photophysiological parameters are described in Supplemental Table 1.

Using a modified version of the absorbance algorithm (Oxborough et al., 2012), volume-based productivity rates (i.e. mol electrons (RCII)^-1^ m^-3^ d^-1^) were calculated as:

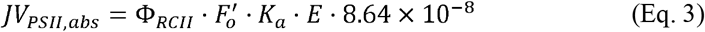

where 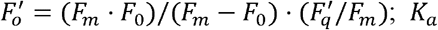 is the instrument calibration factor (11800 m^-1^); E = irradiance (µmol photons m^-2^ s^-1^) of actinic light in the instrument and the factor 8.64 × 10^−8^ converts µmol photons m^-2^ s^-1^ to mol photons m^-2^ d^-1^ and kg to mg. The photon to electron conversion Φ_RCII_ (mol e^-^ mol photon^-1^) has a constant value of 1, representing one electron transferred from P680 to quinone A (Q_A_) for each photon absorbed and delivered to reaction center RCII (Kolber and Falkowski, 1993). RCII was estimated as:

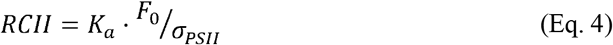

where F_0_ is dark-adapted base fluorescence and σ_*PSII*_ is the absorption cross-section area of the dark-adapted photosystem. Here, we used mean nighttime σ_*PSII*_ measurements and assumed similar absorption cross section during the following day to avoid errors from daytime suppression due to enhanced base fluorescence and non-photochemical quenching. JV_PSII_ (mol electrons m^-3^ d^-1^) was converted to carbon units by the factor Φe:c (Schuback and Tortell, 2019):

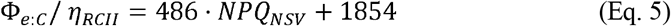

where Φ_e:C_ is the electron to carbon fixation ratio; η_*RCII*_ is the RCII to Chl-*a* ratio; and *NPQ*_*NSV*_ is the normalized Stern-Volmer non-photochemical quenching coefficient. η_*RCII*_ was determined using the RCII calculated in Eq. 4 and dividing by Chl*a* concentration from the ship’s fluorometer. During the timespan where the fluorometer readings were faulty, we used nighttime F_0_ from the FRRf and the relationship between F_0_ and Chl*a* before the sensor was fouled.

To calculate water column GPP from FRRf measurements, we used *in-situ* light attenuation from noon CTD profiles to calculate the light field over the diurnal cycle. P vs. E relationships were determined from FRRf productivity parameters according to Platt et al. (1980) if photoinhibition was observed (Eq. 6) or from Webb et al. (1974) if no photoinhibition was observed (Eq. 7) :

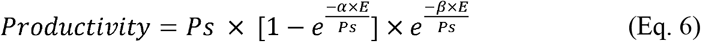

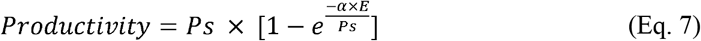

where *Ps* is the maximum photosynthesis, E is PAR, *α* is the initial photosynthesis slope under low irradiance, and *β* is the slope under high/stressful irradiance.

Due to system oligotrophy, some measurements did not pass quality control even at lower light (250 µmol photons m^-2^ s^-1^). These P vs. E curves were omitted from further analysis, but the parameters were used for mean estimates of basic dark adapted photophysiological parameters (see below). Due to instrument failures and missing data, we obtained a higher temporal resolution of GPP by averaging the valid daytime productivity estimates (Eq. 6, 7) for every 30 min at their respective light conditions. We considered the parameters only valid to 30 m depth because cellular Chl*a* changed below 30 m (Supplemental Fig. 3). Additional photophysiological parameters derived from dark-adapted measurements are described in Supplemental Table S1.

### 2.6. Statistical methods

All statistical analyses were conducted in R (version 2024.12.0+467; R Core Team). For comparisons of productivity metrics across cycles and treatments, we assumed normality and homogeneity. Significant differences among Lagrangian cycles were tested by one-way ANOVA followed by post-hoc pairwise comparisons by Tukey’s Honest Significant Difference (HSD) test. Depth-integrated rates (e.g., ^14^C-NPP, GPP, N_2_ fixation) were calculated by trapezoidal integration of depth profiles over the upper 30 m and the full photic zone, with variability reported as standard error of the means (SEM) of replicate profiles. All plots were generated using ggplot2, and compositing of multi-panel figures was performed using patchwork packages. The significance level was set at α = 0.05 for all tests.

## 3. Results

### 3.1. Environmental properties

During the RR2201 cruise, the Argo Basin was characterized by warm surface waters and strong thermal stratification (Fig. 1A, Table 1). Sea surface temperatures (upper 30 m) ranged from ∼28°C to 30°C across all cycles. Subsurface temperatures at the DCM (67–75 m) were ∼4°C lower compared to surface values, highlighting a strong thermal stratification. Salinity remained relatively uniform throughout the surface layer in Cycles 1, 2, and 4 (hereafter C1 to C4), with modest increases at depth (Fig. 1 B). The highest salinity was observed during Cycle 2, while Cycle 3 showed a distinct salinity anomaly near 50□;m depth, potentially indicating a subsurface intrusion of water from lateral advection or a remnant signal of a prior mixing event. Mixed layer depth (Fig. S2, Table 1), based on sigma theta was determined to average around 30.5 ± 3.1, 9.4 ± 1.0, 6.6 ± 0.2 and 9.2 ± 0.8 for C1 to C4, respectively. Mixed layer depth, based on the oxygen profiles used for analysis of the O_2_/Ar data was determined to be 53 ± 7, 32 ± 7, 17 ± 2.5, 20.5 ± 7 m, for C1 to C4, respectively (Fig. S2, Table 1). The generally deeper mixing during C1 was the result of a storm system which passed through the area shortly before C1. Since C3 was the same water mass as C1, re-stratification occurred within 7 days. The alternative mixed layer depth based on the oxygen profile showed consistently deeper values compared to the density-based MLD, particularly during stratified periods. These discrepancies reflect the differing time scales of physical mixing (hours to days) and biological and air-sea gas exchange processes (days to weeks). The latter MLDs are more appropriate for use in the O_2_/Ar NCP assessment.

**Table 1:** Nutrient and biophysiochemical parameters during the cyles, averaged by depth intervals. Where shown, the errors are 1 standard error of the mean.

|  | Depth | Cycle 1 | Cycle 2 | Cycle 3 | Cycle 4 |
| --- | --- | --- | --- | --- | --- |
| Starting Dates |  | 02/04 | 02/09 | 02/15 | 02/20 |
| Par | 0-30 m | 75.80 ± 4.15 | 161.25 ± 7.37 | 156.61 ± 7.52 | 102.63 ± 5.38 |
|  | 30-50 m | 24.42 ± 1.37 | 53.46 ± 2.85 | 53.77 ± 3.05 | 48.39 ± 2.86 |
|  | 50-100 m | 6.35 ± 0.28 | 11.52 ± 0.54 | 10.44 ± 0.52 | 11.84 ± 0.61 |
|  | Deep | 0.17 | 0.17 ± 0.01 | 0.15 | 0.13 ± 0.01 |
| Salinity | 0-30 m | 34.54 | 34.85 | 34.57 | 34.54 |
|  | 30-50 m | 34.54 | 34.89 | 34.71 | 34.54 |
|  | 50-100 m | 34.48 | 34.83 | 34.62 | 34.60 |
|  | Deep | 34.69 | 35.02 | 34.82 | 34.78 |
| Temp | 0-30 m | 28.57 | 28.41 ± 0.01 | 29.08 ± 0.01 | 29.78 ± 0.01 |
|  | 30-50 m | 28.49 ± 0.01 | 27.35 ± 0.03 | 27.87 ± 0.01 | 28.80 ± 0.02 |
|  | 50-100 m | 26.06 ± 0.03 | 24.53 ± 0.02 | 25.40 ± 0.02 | 25.73 ± 0.02 |
|  | Deep | 12.96 ± 0.08 | 16.12 ± 0.06 | 14.45 ± 0.08 | 14.49 ± 0.07 |
| MLD st |  | 41 ± 15 | 10 ± 5 | 6.8 ± 1.2 | 10.76 ± 3.7 |
| MLDO<sub>2</sub> |  | 53 ± 7 | 32 ± 7 | 17 ± 2.5 | 20.5 ± 7 |
| NO<sub>3</sub><sup>-</sup> | 0-30 m | 0.03 ± 0.01 | 0.02 | 0.01 | 0.01 |
|  | 30-50 m | 0.01 | 0.02 ± 0.01 | 0.05 ± 0.03 | 0.01 |
|  | 50-100 m | 1.94 ± 0.75 | 1.33 ± 0.45 | 1.14 ± 0.46 | 2.10 ± 0.91 |
|  | Deep | 21.03 ± 3.90 | 11.92 ± 2.22 | 20.02 ± 3.04 | 13.56 ± 1.83 |
| NH<sub>4</sub><sup>+</sup> | 0-30 m | 0.01 | 0.01 | 0.01 | 0.01 |
|  | 30-50 m | 0.01 ± 0.01 | 0.01 | 0.02 | 0.01 |
|  | 50-100 m | 0.02 | 0.02 | 0.02 | 0.01 |
|  | Deep | 0.01 | 0.01 | 0.01 | 0.01 |
| PO<sub>4</sub><sup>3-</sup> | 0-30 m | 0.11 ± 0.01 | 0.05 ± 0.01 | 0.05 | 0.06 ± 0.02 |
|  | 30-50 m | 0.10 ± 0.01 | 0.04 | 0.06 ± 0.01 | 0.08 ± 0.02 |
|  | 50-100 m | 0.24 ± 0.05 | 0.20 ± 0.03 | 0.20 ± 0.04 | 0.30 ± 0.06 |
|  | Deep | 1.55 ± 0.27 | 0.85 ± 0.14 | 1.45 ± 0.21 | 0.95 ± 0.12 |
| NO<sub>2</sub><sup>-</sup> | 0-30 m | ND | ND | ND | ND |
|  | 30-50 m | ND | ND | ND | ND |
|  | 50-100 m | 0.07 ± 0.02 | 0.08 ± 0.03 | 0.07 ± 0.03 | 0.07 ± 0.02 |
|  | Deep | 0.03 ± 0.01 | 0.02 | 0.01 ± 0.01 | 0.01 ± 0.01 |
| SiO<sub>4</sub> | 0-30 m | 2.00 ± 0.22 | 1.85 ± 0.02 | 1.79 ± 0.04 | 1.77 ± 0.02 |
|  | 30-50 m | 1.80 ± 0.08 | 1.97 ± 0.05 | 2.13 ± 0.08 | 2.02 ± 0.39 |
|  | 50-100 m | 3.85 ± 0.70 | 3.71 ± 0.33 | 4.13 ± 0.48 | 5.00 ± 0.77 |
|  | Deep | 44.31 ± 13.22 | 17.37 ± 5.31 | 40.23 ± 10.29 | 16.16 ± 2.81 |

Nutrient concentration profiles verified the oligotrophic nature of the basin (Fig. 2, Table 1). Nitrate concentrations (NO_3_^-^) were generally depleted (Fig. 2F), averaging (± SEM) 0.018 ± 0.04 µM in surface waters down to 50 m, with a nitracline (defined as the depth at which nitrate concentrations first exceed 1 µmol L^-1^) around 65-81 m. Phosphorus (Fig. 2H), showed a residual concentration of 0.07 ± 0.02 µM PO_4_^3-^ µM. P* (the overabundance of phosphorus relative to nitrogen P* = [PO_4_^3-^] – 16/[NO_3_^-^]) averaged 0.06 in the upper 50 m and 0.1 for the upper 100 m (Figure Supplemental 2), indicating that nitrate was a limiting major macro-nutrient in the surface ocean. Silicic acid (Fig. 1H) was measured at average concentrations of 1.89 ± 0.04 µM within the upper 50 m; ammonia (NH_4_^+^) was measurable (0.012 1 µM) throughout the water column (Fig. 1I), indicating active recycling. Nitrite (Fig. 1G) had a prominent maximum, indicating enhanced nitrification, slightly below the deep chlorophyl maximum (DCM) (Fig. 1E). The DCM (Fig. 1E) depths ranged from 67 to 75 m, with the nitracline typically slightly deeper than the DCM. The euphotic zone (EZ) depth (1% surface PAR) (Fig. 1D) was slightly deeper than the DCM, while subsurface fluorescence peaks marked the DCM and associated productivity zones.

**Fig. 2.**
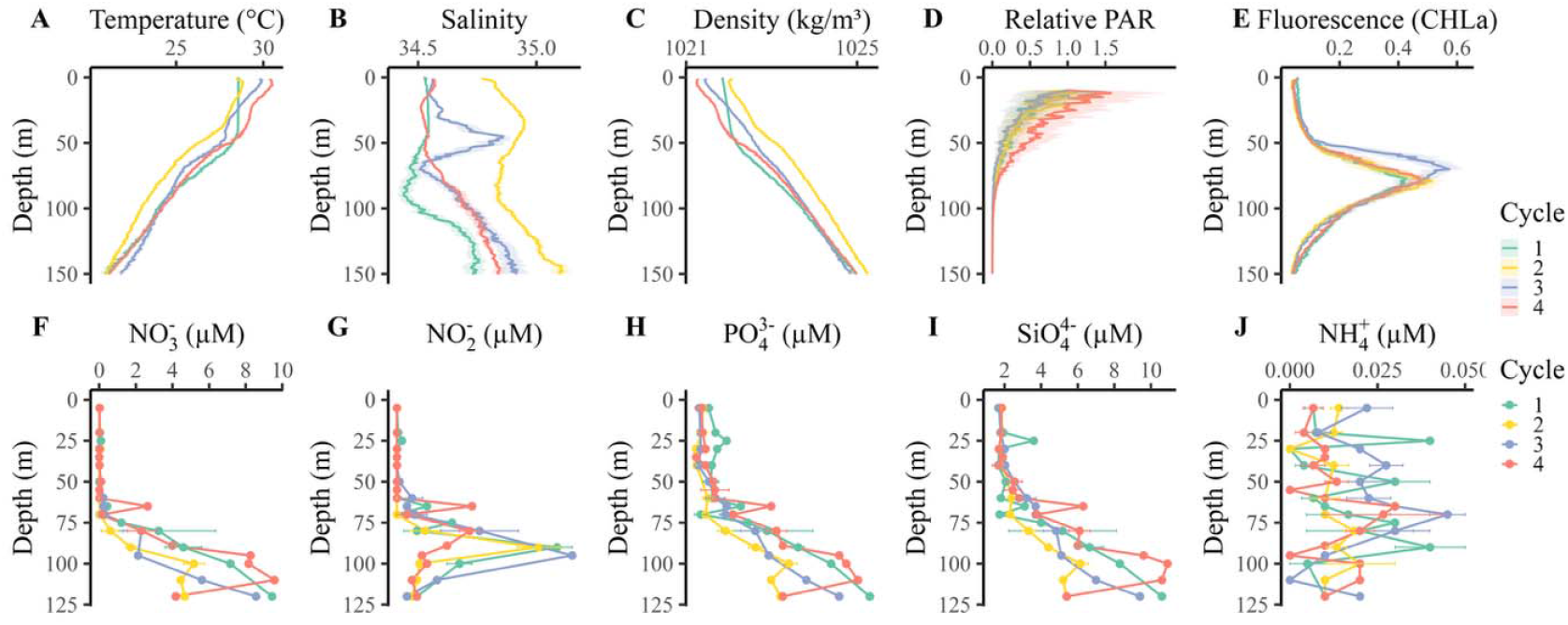
Physical and biochemical and profiles from the 4 cycles plotted to 150-m depth. A) Temperature (°C), B) Salinity (PSU), C) Density (kg m^-3^), D) Relative PAR (µmol photons m^-2^ s^-1^), E) Chl*a* fluorescence (µg L^-1^) taken from the CTD rosette depth profiles and plotted as average per cycle with 95% confidence interval (shaded area). F-J) Nutrient profiles (µM) taken from CTD niskin bottles. F) NO_3_^-^, G) NO_2_^-^, H) PO_4_^3-^, I) SiO_4_^4-^ J) NH_4_^+^. Error bars denote standard deviation when multiple samples were taken from the same depth during a cycle.

**Fig. 3.**
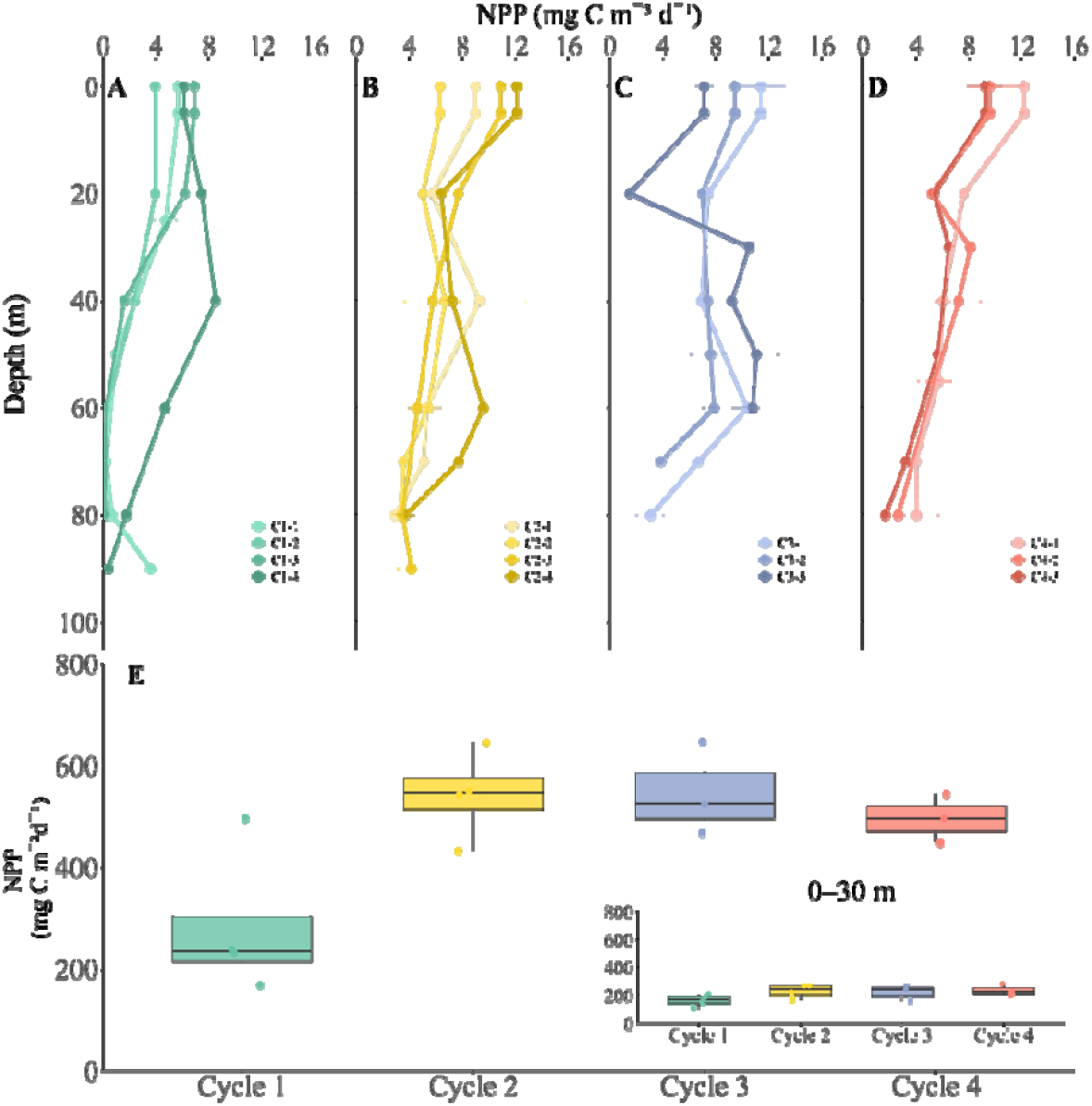
Daily measurement profiles and integrated values of ^14^C based net primary productivity (NPP_14C_). A-D) NPP_14C_ profiles for all cycles and cycle days with error bars denoting 1 standard error of the mean. E) Integrated productivity in mg C m^-2^ d^-1^. Dots indicate the water column productivity for each day (n = 4, 4, 3, and 3 for Cycles 1, 2, 3, and 4, respectively) while the boxplot indicates the median and interquartile range of integrated values. Insert: Depth integrated NPP_14C_ of the water column of the upper 30 m.

**Fig. 4.**
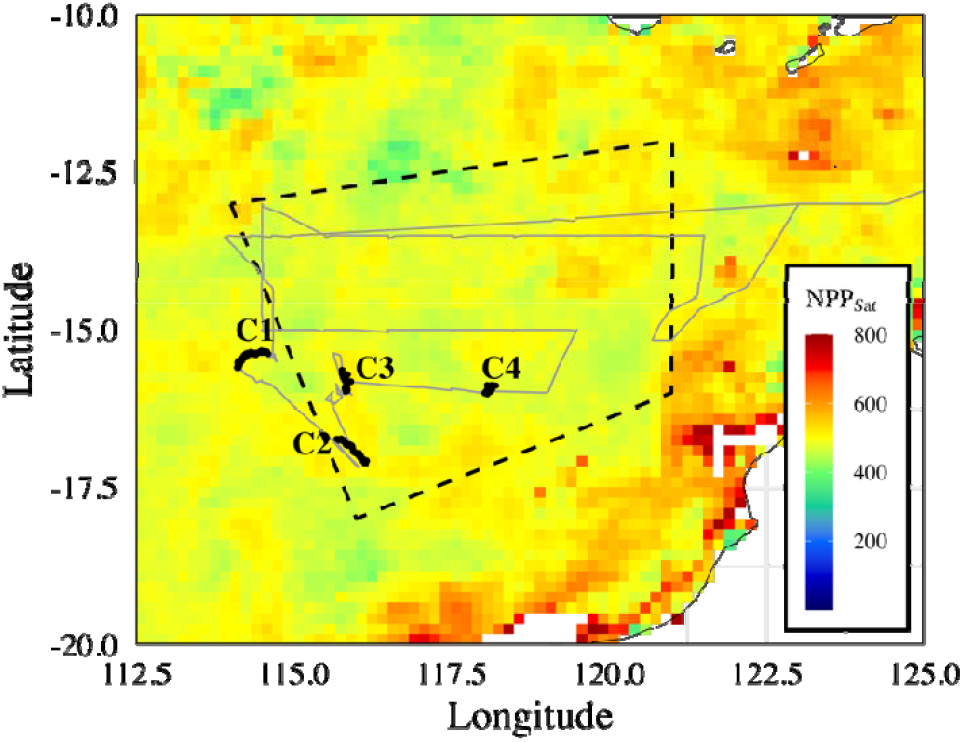
Satellite observed net primary productivity (NPP_Sat_). The underlying data are from the CAFÉ model extracted for Feb. 10-17. The cruise track is shown in gray, the different cycles are indicated in black and the polygon indicated the study region as analyzed by Kehinde et al. (2022) bounded by coordinates: 13°S, 114°E; 18°S, 116°E; 16°S, 121°E; 12°S, 121°E).

#### 3.2. Net primary productivity (NPP) on in-situ array

^14^C-based NPP profiles showed distinct differences across the four Lagrangian experiments. For C1, NPP_14C_ was greatly reduced in the lower euphotic zone (EZ) due to low light (cloud cover) during the first 3 days (Fig. 2), but increased sharply on day 4. For C2 and C3, profiles were relatively constant to 60 m. In contrast, C4 showed a strong, almost linear, decrease with depth, which correlated with increasing cellular Chl*a* below 40 m (Fig. S3).

Integrated NPP_14C_ rates were relatively consistent across the cycles except for days 1-3 of C1. Cycle-averaged productivity was lowest in C1 (283 ± 72 mg C m^-2^ d^-1^), and C2-C4 exhibited higher full depth integrated NPP_14C_ values of 542 ± 43, 546 ± 53, and 496 ± 28 mg C m^-2^ d^-1^, respectively. The overall mean NPP_14C_ across all four cycles was 460 ± 40 mg C m^-2^ d^-1^. Integrated ^14^C-based primary productivity for the upper 30 m (the habitat depth of Southern Bluefin Tuna larvae) (Fig. 2E insert, Table 2) averaged roughly half of the full EZ-integrated rates. Cycle mean values ranged from 164 ± 20 mg C m^-2^ d^-1^ in C1 to 240 ± 21 mg C m^-2^ d^-1^ in C4, with an overall average of 212 ± 14 mg C m^-2^ d^-1^.

**Table 2:**
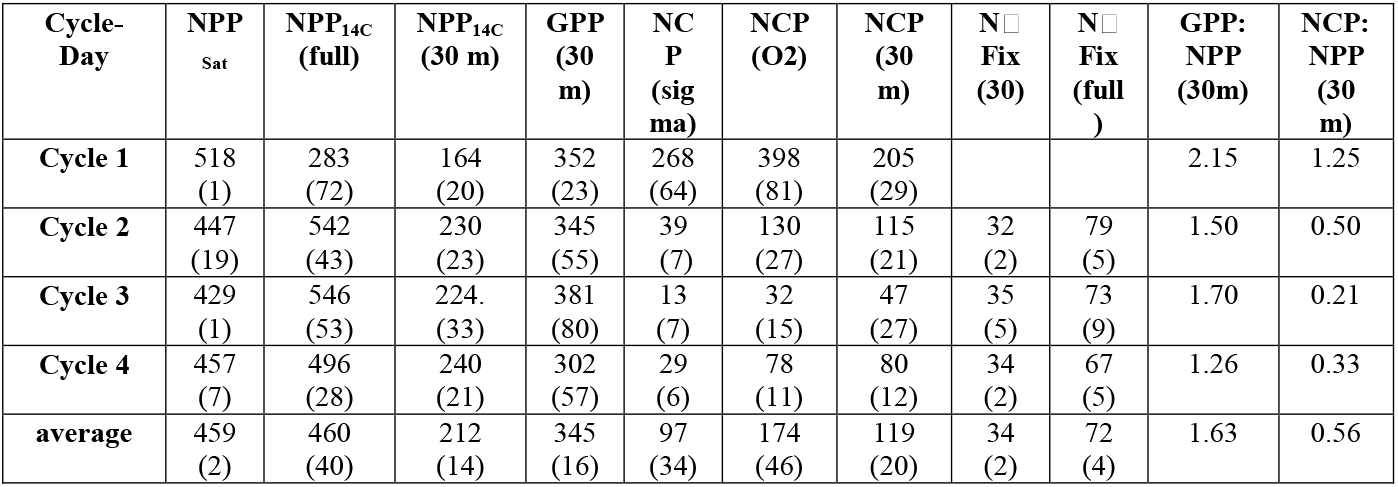
Rates of productivity measurements and ratios. Only average values are presented in this table. Cycle day values are presented in supplemental Table S2. Rates are expressed in mg C m^-2^ d^-1^. Numbers in parenthesis denote standard error of the mean.

Supplementing our direct NPP_14C_ measurements, NPP_Sat_ estimates for the region averaged 450 mg C m^-2^ d^-1^ along the cruise track (Fig. 3). Productivity was highest, up to a maximum of ∼900 mg C m^-2^ d^-1^, in patches in the shelf area between 120 and 125°E and along the Australian coastline south of Argo Basin. West of 120 °E, productivity between 12.5 and 17.5°S was relatively uniform (464 mg C m^-2^ d^-1^ on average), showing no obvious indications of local production hot spots in the Argo Basin during our cruise (Fig. 3).

For the Lagrangian cycles, 8-day NPP_Sat_ estimates ranged from 428 to 518 mg C m^-2^ d^-1^ (Table 2), with an overall cycle mean of 459 ± 2 mg C m^-2^. Post□;hoc Tukey’s HSD tests indicated significant differences among cycles. During C1, mean NPP_Sat_ (515 mg C m^-2^ d^-1^) was significantly higher than C2-C4 (p < 0.01). In C2, productivity was lowest (446 ± 19 mg C m^-2^ d^-1^; p=0.002 versus C1) and exhibited the highest day-to-day variability. NPP_Sat_ for C3 (Feb. 19– 26; 429 ± 1 mg C m^-2^ d^-1^) was significantly lower than both C1 (p<0.001) and C4 (459 ± 2 mg C m^-2^ d^-1^; p=0.03).

#### 3.3 N_2_ fixation on in-situ array

N_2_-fixation rates are unavailable for C1 and the first two days of C2 due to lost samples. Measured rates ranged generally from 10 to 20 nmol N_2_ L^-1^ d^-1^ with an average of ∼15 nmol N_2_ L^-1^ d^-1^ for the upper 60 m, decreasing sharply to 1-2 nmol N_2_ L^-1^ d^-1^ at depths of 70-80 m in the nitracline. Based on Redfield stoichiometric conversion to mg C m^-2^ d^-1^, EZ-integrated rates ranged from 60 to 86 mg C m^-2^ d^-1^, with the highest value on day 3 of C3 (84 ± 4 mg C m^-2^ d^-1^) and the lowest on day 3 of C4 (57 ± 3 mg C m^-2^ d^-1^) (Fig. 5, Table 2). Corresponding 30-m integrated rates ranged from 27 ± 6 to 41 ± 0 mg C m^-2^ d^-1^, averaging 34 ± 2 mg C m^-2^ d^-1^. N_2_-fixation for the full depth was highest during C2 (84 ± 3 mg C m^-2^ d^-1^), followed by C3 (76 ± 7 mg C m^-2^ d^-1^) and C4 (61 ± 4 mg C m^-2^ d^-1^), giving an overall average of 74 ± 3 mg C m^-2^ d^-1^ (Table 2).

**Fig. 5:**
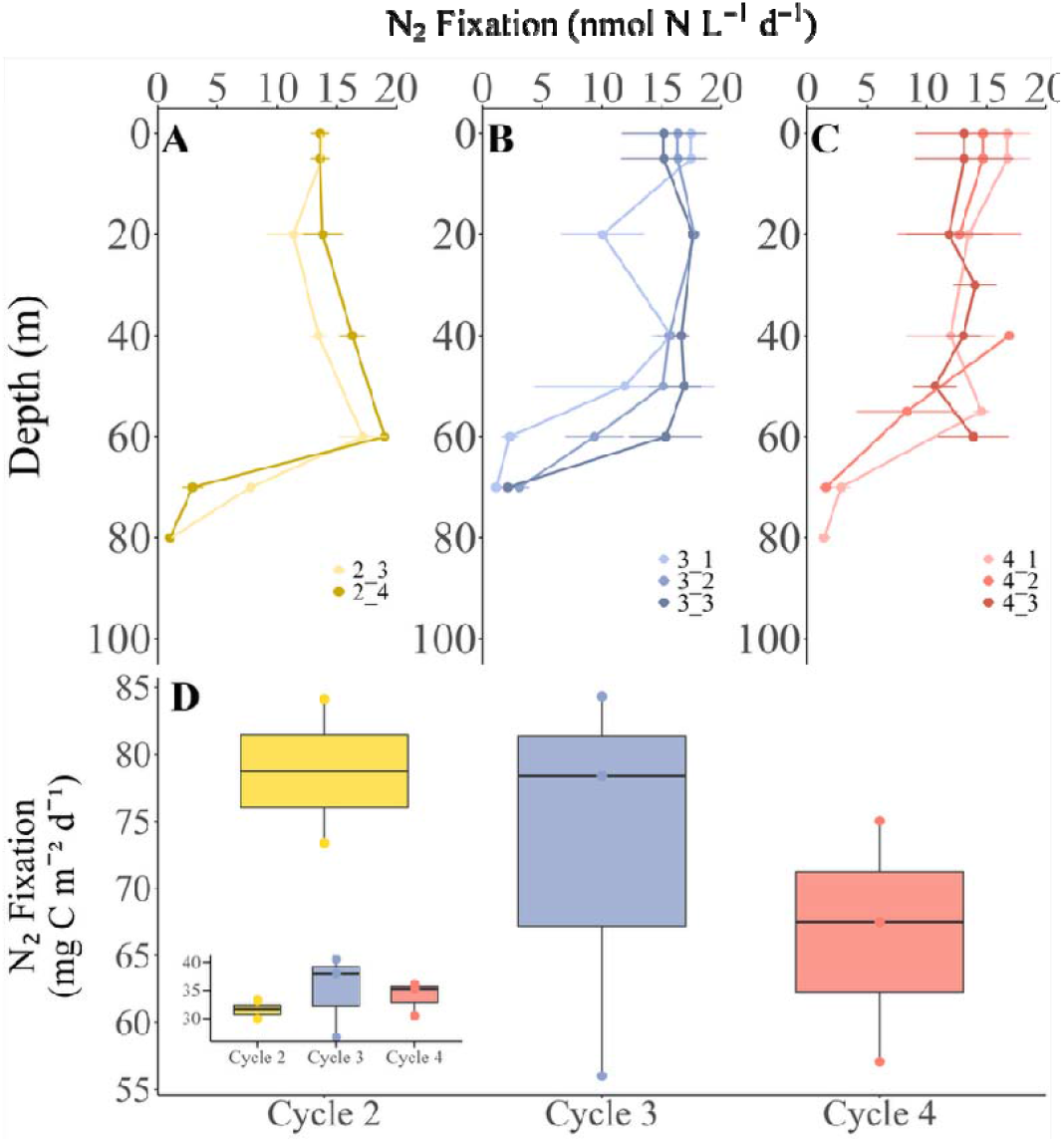
N_2_ fixation profiles and integrated rates. A-D); N_2_-fixation over the depth of the array in nmol N m^-3^ d^-1^ for all cycles and cycle days. D) Integrated N_2_-fixation in mg C m^-2^ d^-1^. Dots indicate the water column integrated productivity for each day (n = 2, 3, 3 for Cycles 2, 3, and 4, respectively), while the boxplot indicates the median and interquartile range. Insert: Depth integrated N_2_ fixation of the water column of the upper 30 m.

#### 3.4. Gross primary productivity (GPP)

Due to low biomass, about 30% of the P vs E curves did not pass quality control (i.e. low signal-to-noise ratio resulted in low data confidence at light levels >250 µmol photons m^-2^ s^-1^). Because community compositions (Selph et al., this issue; Yingling et al., this issue) and base GPP calculation parameters were similar among cycles (Figure S3), we used the diurnal average of the fitted photosynthetic parameters (α, P_max_, β) from the P vs. Es equations to substitute missing data (see methods, Eq. 6, 7). These data (Fig. 6 A-D), revealed a midday productivity saturation for most cycles (e.g., flat production despite the sinusoidal solar cycle).

**Fig. 6:**
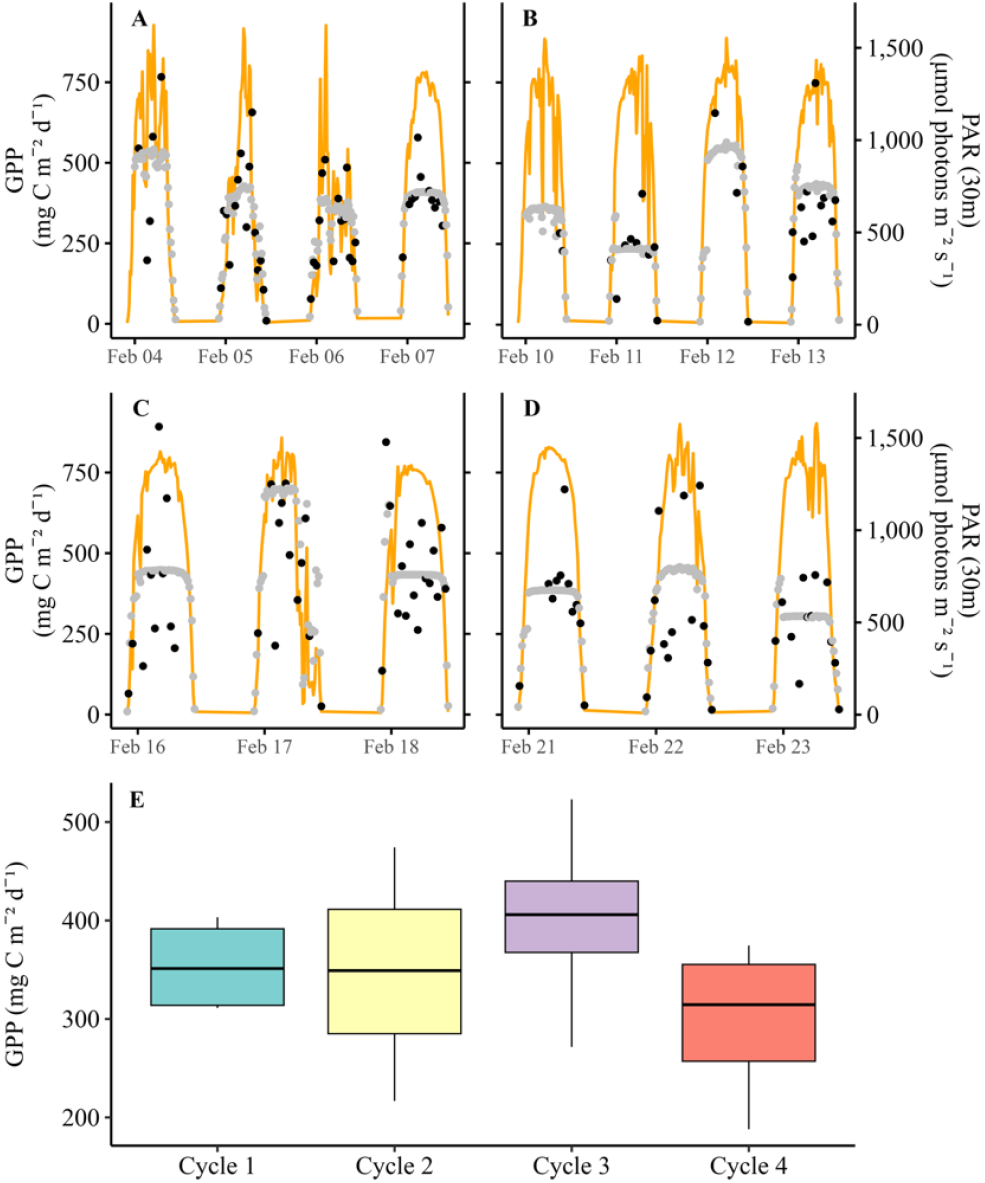
Gross primary productivity over the diurnal cycles and integrated over 30 m. Panels A-D correspond to C1-C4, respectively. Orange lines indicate the median light intensity for 30 m water depth in µmol photons m^-2^ s^-1^. Black dots indicate the calculated GPP for all valid FRRf P vs. E curves integrated over the upper 30 m. Grey dots indicate the interpolated data based on 30 min intervals using the average P vs. E (alpha, Pmax, beta) integrated over the upper 30 m. D: Average GPP for each cycle.

FRRf-based estimates of gross primary productivity for the upper 30 m (GPP_30m_) varied daily from 188 to 465 mg C m^-2^ d^-1^, with a cycle average of 345 ± 16 mg C m^-2^ d^-1^ (Fig. 6E, Table 2, Table S2). Despite reduced PAR at the beginning, C1 showed similar GPP rates compared to the other cycles (mean 352 ± 23 mg C m^-2^ d^-1^), with daily values from 305 to 403 mg C m^-2^ d^-1^ (Table S2). C2 showed comparable day-to-day variability (217 to 465 mg C m^-2^ d^-1^) and a similar mean value (345 ± 55 mg C m^-2^ d^-1^). C3 had the highest mean GPP_30m_ at 381 ± 80 mg C m^-2^ d^-1^, peaking at over 500 mg C m^-2^ d^-1^ on day 3. In contrast, C4 displayed the lowest GPP_30m_, ranging from 188 to 375 mg C m^-2^ d^-1^ with a cycle mean of 302 ± 57 mg C m^-2^ d^-1^. Despite these cycle differences, a one-way ANOVA using daily GPP_30m_ revealed no statistically significant differences among cycles (*p* > 0.05). Direct comparison of GPP_30m_ to NPP_14C_ for the upper 30 m resulted in a mean gross-to-net productivity ratio (GPP:NPP_30m_) of 1.6 (Table 2) with high variability in individual daily estimates (0.67 to 3.21 (Table S2).

#### 3.5. Net community productivity (NCP)

Net community productivity was calculated based on O_2_:Ar data, wind velocity and mixed layer depth. MLD_σθ_ estimates for C2 to C4 were below 10 m, while O_2_ profiles indicated a deeper MLD (Fig. S2). Hence, we analyzed NCP based on both MLD_σθ_ and MLD_O2_ (hereafter, NCP_σθ_ and NCP_O2_, respectively). In addition, we calculated a NCP_30m_ estimate based on a fixed 30-m MLD for direct comparisons with GPP_30m_ and NPP_30m_.

The relatively shallow MLD_σθ_ resulted in short O_2_ residence time; hence, daily changes in wind affected NCP calculations most strongly (Fig. 7). C1 had the highest NCP_σθ_ (Table S2), ranging from 116 to 388 mg C m^-2^ d^-1^ (Table 2), with a cycle mean of 268 ± 64 mg C m^-2^ d^-1^. This is likely due to the mixing from the previous storm event as well as the integration over a deeper mixed layer. NCP_O2_ and NCP_30m_ estimates for C1 were 398 ± 81 and 205 ± 29 mg C m^-2^ d^-1^, respectively (Table 2, Fig. 7). C2 exhibited substantially lower NCP_σθ_, NCP_O2_ and NCP_30m_ estimates of 39 ± 7, 130 ± 27 and 115 ± 21 mg C m^-2^ d^-1^, respectively, reflecting the shallower MLD. C3 displayed the lowest values of 13 ± 7 (NCP_σθ_), 32 ± 15 (NCP_O2_), and 47 ± 27 mg C m^-2^ d^-1^ (NCP_30m_). In C4, the respective estimates were 29 ± 6, 78 ± 11, and 80 ± 12 mg C m^-2^ d^-1^.

**Fig. 7.**
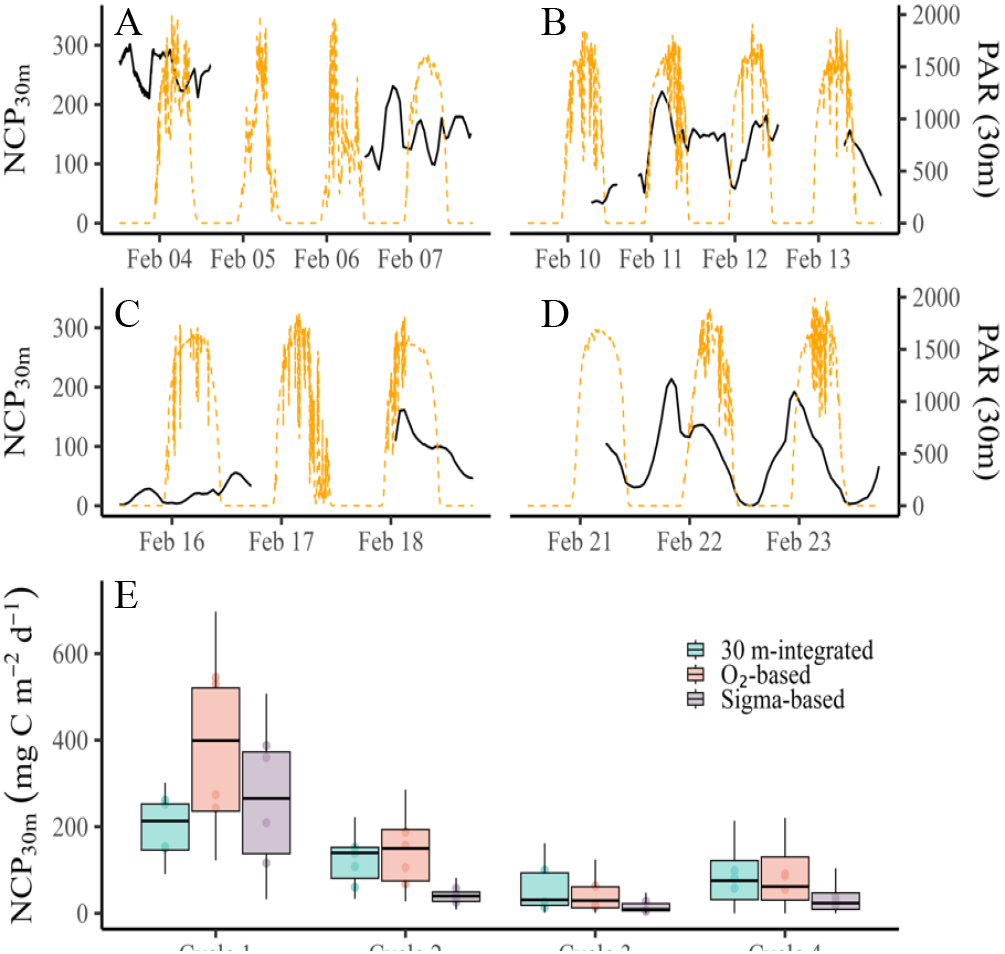
Net Community Production (NCP, mg C m^-2^ d^-1^) in each cycle. Panels A-D: Solar irradiance in orange; NCP calculated using MLD_30m_ in black over the day-night times for each cycle. E) NCP averaged over the cycle duration: Box plot indicates the calculations based on the different MLD applied (30-m integrated, O_2_-based, or Sigma-based).

Across all cycles, NCP_30m_ and NPP_30m_ averaged 119 ± 20 and 212 ± 14 mg C m^-2^ d^-1^, respectively, resulting in a mean NCP:NPP_30m_ ratio of ∼0.56 (Table 1). These results suggest that just over half of net primary production was retained within the upper 30 m, with the remainder lost to community respiration or export. The lowest retention NCP:NPP_30m_ = 0.21 occurred in C3, highlighting some temporal variability in ecosystem metabolic balance.

#### 3.6. Photophysiology

During C1, lower light in the upper 30 m (PAR_30m_ = 452 ± 8 µmol photons m^-2^ s^-1^) yielded a moderate PSII efficiency (Fv/Fm= 0.26 ± 0.002) and a light-saturation irradiance of 334 ± 9 µmol photons m^-2^ s^-1^ (Fig. S3). Under these lower□light conditions, cells showed a larger antenna size (σ = 6.62 ± 0.07 nm^2^), significantly greater than C2 and C3 (p < 0.01), and exhibited modest non photochemical quenching in the dark-adapted state (NSV = 2.89 ± 0.01). Electron transport through the photosystem II (1/τ□□= 1.25 ± 0.04 ms^-1^) was moderate (Fig. S3). For C2, average light intensity (PAR_30m_) was 598 ± 9 µmol photons m^-2^ s^-1^ with a low photosynthetic quantum yield (Fv/Fm = 0.19 ± 0.003, the lowest of all cycles (p < 0.0001) and high Ek (412 µmol photons m^-2^ s^-1^). Antenna size was significantly reduced compared to C1 (σ = 5.08 ± 0.11 nm^2^; p < 0.001), while non-photochemical quenching in the dark-adapted state increased (4.52 ± 0.10; p < 0.01) (Fig. S3). The electron transport rate through the photosystem (1/τ) increased slightly (1.4 ± 0.1 ms^-1^), consistent with accelerated electron turnover under photo-stress. C3, the continuation of C1, exhibited similar mean light intensity compared to C2 (559 ± 10 µmol photons m^-2^ s^-1^), and showed elevated photochemical efficiency (Fv/Fm= 0.30 ± 0.002; p < 0.01 versus C1) and light□;use efficiency at sub□;saturating irradiance (α = 0.046 ± 0.001; p < 0.001) and lowest Ek (283 ± 10; p < 0.01 µmol photons m^-2^ s^-1^ versus C1 and C2). Both σ (5.84 ± 0.04 nm^2^) and nonphotochemical quenching (2.40 ± 0.03) were low, and 1/τ□;□; was lowest (0.95 ± 0.03 ms□;^1^; p < 0.01) (Fig. S3)., indicating optimal light capture and minimal energy dissipation under balanced light conditions. During C4, the average light intensity (PAR30m = 579 ± 10 µmol photons m^-2^ s^-1^) was similar to C2 and C3, with moderate PSII efficiency (Fv/Fm = 0.23 ± 0.003). The light-saturation irradiance (Ek = 302 ± 14 µmol photons m^-2^ s^-1^) was intermediate between C1 and C3. Antenna size (σ = 6.17 ± 0.08 nm^1^) showed a moderate value, closer to that of C1, while non-photochemical quenching (NSV = 3.36 ± 0.06) indicated a moderate energy dissipation capacity. Electron transport rate through the photosystem (1/τ□□; = 1.40 ± 0.04 ms^-1^) was similar to C2, suggesting comparable electron turnover rates under these intermediate conditions (Fig. S3).

## 4. Discussion

### 4.1. General features of the Argo Basin

Our study revealed warm surface temperature, a mostly shallow MLD and near-complete depletion of nitrate in surface waters of the Argo Basin (Fig. 1). Residual phosphorus of 0.06 µmol L^-1^ gave ratios of dissolved inorganic N:P well below Redfield. The nitracline was consistently between 70 to 81 m, similar to the 67–75 m DCM (Fig. 2E). Residual ammonium (∼0.01 µmol L^-1^; Fig. 2J) throughout the EZ was consistent with efficient N recycling. Residual silicate (Fig. 2I; ∼1.9 µmol L^-1^) and available P aligned with dominance of small cyanobacteria such as *Prochlorococcus* and low abundances of silicified phytoplankton, like diatoms (Yingling et al., this issue). Additionally, the deep nitrite maxima (Fig. 2G) and O_2_ depletion near the DCM (Figure S2; (Stukel et al., this issue) suggested enhanced nitrification and microbial respiration, potentially supporting microaerobic processes such as denitrification or anammox (Dore et al., 2002; Fawcett et al., 2015). Despite these nutrient features (Fig. 2) and high cellular Chl*a* in the lower EZ (Fig. S1; (Stukel et al., this issue), >50% of total NPP occurred in the upper 50 m (Fig. 2), indicating light limitation at depth (Geider et al., 1997; Kana et al., 1997). This was also notable as cloud-suppressed NPP_14C_ during the first three days of C1 (Fig. 3A) and as substantially reduced phytoplankton growth rates from dilution experiments incubated in the lower EZ (Landry et al., this issue).

Together, these features illustrate a recycling-dominated system, where N availability restricts surface productivity, light limits sub-surface productivity and recycling processes sustain much of the community’s N demand. The persistent low N:P ratios suggest favorable conditions for diazotrophy (Landolfi et al., 2018; Louchard et al., 2023), although the measured N_2_ fixation rates (Fig. 5) and residual P suggest that diazotroph potential might not have been fully realized during our study period.

### 4.2. Net primary productivity

Our observed rates of NPP_14C_ are broadly consistent with long-term averages from oligotrophic gyres such as HOT and BATS (Brix et al., 2006), but 40% higher than the mean estimate of 325 mg C m^-2^ d^-1^ for the Atlantic Bluefin Tuna spawning region in the Gulf of Mexico (Yingling et al., 2022). Productivity was highest in the upper 50 m, with subsurface productivity following decreasing light availability. The light-limiting nature of subsurface production was further supported by flow cytometric measurements showing increased cellular Chl*a* in *Prochlorococcus* and *Synechococcus* below 40 m (Fig. S1), as well as increases in Chl*a*:POC ratios below 50 m (Selph et al., this issue; Stukel et al., this issue). This pattern parallels previous observations from oligotrophic regions where DCMs are largely driven by pigment adjustments to low-light conditions (Mignot et al., 2014; Cornec et al., 2021; Phongphattarawat et al., 2023).

Comparing *in-situ* daily productivity (NPP_14C_) to remotely sensed estimates using the 8-day composite CAFE model (NPP_Sat_), NPP rates converged on similar cycle-averaged values of 460 mg C m^-2^ d^-1^ (Figs. 2 and 3, Table 1), supporting the general usage of such algorithms to assess productivity of this region. However, substantial discrepancies emerged on shorter timescales (Table S2, Fig. 3, 5, 6). For example, the day-to-day variability of NPP_14C_ in C1 (Fig. 2A) was not captured by NPP_Sat_ (Table S2), which remained stable throughout the observational period due to the longer time integration. Variability in NPP_14C_ reflects transient physical or biological processes (e.g., short-term mixing, changes in light availability and photoacclimation). While satellite algorithms have made progress in capturing depth-dependent distribution of biomass, Chl*a* and estimated productivity (Westberry et al., 2008), the differences in our study are likely driven by their limited ability to resolve short-term changes in subsurface production in stratified deep oligotrophic systems.

### 4.5. Support of NPP by N_2_ fixation

N_2_ fixation was estimated to be a secondary N source supporting primary production. As shown in Yingling et al. (this issue) ammonium utilization was the primary N source for the community while nitrate uptake was mostly negligible in surface waters. Using Redfield stoichiometry for C:N conversion, N_2_-fixation rates for C2–C4 averaged 34 ± 2 mg C m^-2^ d^-1^ in the upper 30 m and 72 ± 4 mg C m^-2^ d^-1^ for the full profile (Fig. 5, Table 2) supporting16% of measured NPP_14C_. Hence N_2_ fixation presents a substantial contributed to the “new” N pool in this region (Eppley and Peterson, 1979). More importantly for the biogeochemistry of this region, the N_2_ fixation rates measured account for the majority of new N required to balance the measured carbon export of 19% of NPP_14C_ into sediment traps (Stukel et al., this issue).

Complementary on-deck incubations performed on the cross-basin transects following C4 revealed more variable rates further in the central Argo Basin, and diel experiments (12 h daytime) indicated 2-3 fold higher daytime than nighttime rates. These rates would indicate that both light-dependent and light-independent N_2_-fixation pathways contributed to the N budget in this region. Interestingly, residual phosphorus concentrations, low N:P ratios, warm temperature and strong stratification are considered diazotrophy-favorable conditions. Yet, the residual phosphorus also indicate that N_2_ fixation was not fully realized. Ecological controls such as trace metal limitation and/or partial light limitation deeper in the water column may have constrained overall N fixation activity, as previously observed in other parts of the Indo-Pacific (Raes et al., 2015; Landolfi et al., 2018; Chowdhury et al., 2023).

### 4.3. Gross production and autotrophic respiration

Primary productivity involves the harvesting of light to split water, yielding energy and reductant for nutrient and carbon uptake and fixation (Falkowski, 2012). Variable fluorescence measurements quantify electron generation and transport rates through the photosystem. The method applied here is based on single turnover excitement of the photosystem, measured as photosystem II fluorescence (Oxborough et al., 2012; Schuback et al., 2015; Boatman et al., 2019; Kranz et al., 2020). The FRRf data provide estimates of gross primary production expressed in electrons per photosystem II reaction center. Carbon-based GPP estimates derived from FRRf measurements depend strongly on electrons used per carbon fixed (e:C) ratios, which can vary with species composition, light acclimation and nutrient status. In the present study, neither community composition (Yingling et al., this issue) nor nutrient status changed dramatically and light availability was similar in C2 – C4 (but more limiting during C1). Hence, our cycle estimates should be reasonably comparable among themselves, but absolute values might be over- or underestimated (Halsey et al., 2013; Schuback et al., 2015; Boatman et al., 2019). Underway surface analysis from the ship’s seawater intake ties the GPP estimates to the upper EZ, as both cellular pigmentation and photoacclimation dramatically change at greater depth. Based on flow cytometric data (Fig. S1), cellular Chla values were relatively stable to ∼40 m but increased by 4-5 fold in the deeper EZ. GPP_30m_ rates of ∼380 mg C m^-2^ d^-1^ were comparable to estimates for mid-to-offshore waters of the California Current Ecosystem (Kranz et al., 2020) but lower than the 440 and 710 mg C m^-2^ d^-1^ winter/summer estimates for station ALOHA based on oxygen isotope data (Juranek and Quay, 2005). However, because these rates were integrated over a deeper MLD (45-85 m) depth, our data seem to be within a generally expected range.

GPP:NPP_30m_ ratios, using a 30-m integrated NPP_14C_ data subset, ranged between 0.67 and 3.2 with an average of 1.63 (Table 1). A ratio of 1.63 is consistent with prior observations in oligotrophic regions where cellular respiration accounts for 40–70% of gross photosynthesis (Laws et al., 2000; Marra and Barber, 2004; Halsey et al., 2013; Huang et al., 2021) with even higher % of respiration conducted by diazotrophic organisms (Kana, 1993; Milligan et al., 2007; Halsey et al., 2010; Kranz et al., 2010; Grosskopf and LaRoche, 2012). The higher GPP relative to NPP_14C_ highlights the substantial respiratory losses within the phytoplankton community and underscores the metabolic cost of maintaining growth under low-nutrient conditions (Halsey et al., 2010; Kranz et al., 2011; Halsey et al., 2013; Eichner et al., 2014; Halsey et al., 2014). Lower GPP_30m_ estimates compared to NPP_30m_ (observed C3, day 1 and C4, day 1) do, however, highlight some uncertainty of the FRRf method. Nonetheless, these GPP_30m_ estimates, when considered alongside NPP_30m_ and NCP_30m_ measurements, provide important constraints on trophic efficiency and organic carbon cycling in the region.

### 4.4. Community metabolism, new and export production using O□;/Ar analysis

In regions characterized by high regenerative productivity, such as the Argo Basin, the balance between NPP, NCP and export of particulate organic carbon (POC) is often assumed to approach a quasi-steady state where NCP approximates export production if no significant accumulation of organic matter occurs (Williams et al., 2004; Brix et al., 2006; Kranz et al., 2020). In oligotrophic systems, export production primarily reflects rapid sinking of zooplankton fecal pellets, enabling efficient vertical transfer of organic carbon despite generally low productivity (Stukel et al., this issue). This export relies on new production fueled by external N input, either via vertical supply of NO_3_^-^ or biological N_2_ fixation.

Our O_2_:Ar-based NCP estimates were particularly sensitive to assumptions regarding MLD, which was very shallow based on σ_θ_ criteria. The more appropriate MLD assumptions (MLD_O2_ or the average of MLD_30 m_) resulted in mean NCP estimates of 174, and 119 mg C m^-2^ d^-1^, respectively (Fig. 7, Table 2). In the following we primarily utilized NCP_30m_ estimates, to be able to compare water column NCP rates to GPP and NPP measurements, however, all other rates are also listed in Table 2.

The elevated NCP observed during C1 coincided with prior water column mixing, including potential entrainment of oxygen-rich upper DCM subsurface waters (Fig. S2). Elevated NCP compared to e.g. NPP can be observed under certain physical forcings such as in upwelling regions (Kranz et al., 2020; Wang et al., 2020) or as seen here after a mixing event. The low O_2_:Ar-derived NCP rates underscore the recycling-dominated nature of the Argo Basin. Export efficiencies (ratio of export to NPP), derived independently from sediment trap fluxes and NPP estimates, ranged from 0.085 to 0.23 (average: 0.19) across all cycles (Stukel et al., this issue). These export ratios are notably higher than expected for a small-phytoplankton-dominated oligotrophic system (Buesseler and Boyd, 2009; Siegel et al., 2014; Karl et al., 2021; Stukel et al., 2024). Applying the average NPP_14C_ (460 mg C m^-2^ d^-1^) and an export ratio of 0.19 yields an expected NCP of ∼90 mg C m^-2^ d^-1^, closely matching our cycle-averaged NCP_30m_ values (Table 2, Fig. 6, but exceeding the estimates from C2–C4 by a factor of 2–3. The difference of NCP_30m_ compared to export estimates may partially reflect integration mismatches, as sediment trap fluxes were made just beneath the base of the EZ (116 - 127 m), while O_2_/Ar estimates were constrained to a shallower mixed layer.

The persistent positive NCP across the observation period further supports the net autotrophic status of the system, supplying organic matter to higher trophic levels which might be respired below the MLD as indicated by the strong negative O_2_ gradient below the DCM (Supplemental Fig. S2). Since the Argo Basin expressed higher than expected NCP rates it appears that higher trophic levels may effectively transfer energy through microbial and mesozooplankton food webs. This is supported by food-web analysis suggesting enhanced trophic transfer efficiencies (Landry et al., this issue).

### 4.5. Photophysiology

Analysis of the photosynthetic processes of electron generation and transport, energy dissipation and light use efficiency can shed light on effectiveness of potential limitations of the photoautotrophic productivity and the cells’ ability to adjust to the specific environmental condition found in the Argo basin. During C1, reduced light availability following a transient mixing event resulted in moderate photosynthetic efficiency (Fv/Fm = 0.26) (Fig. S3A,B) and a light-saturation point (Ek) of ∼334 µmol photons m^-2^ s^-1^ (Fig. S3 C). The phytoplankton community responded by increasing absorption cross section (σ = 6.6) (Fig. S3 E), indicative of a larger light-harvesting antennae to optimize photon capture under low light (MacIntyre et al., 2000; Falkowski and Raven, 2007). Non-photochemical quenching (NSV) remained modest (Fig. S3 F), indicating limited excess energy dissipation, while moderate electron turnover rates (1/τ) (Fig. S3 G) suggested balanced excitation pressure and effective electron transport through the lower photosystem (Ruban, 2016). These photoacclimation responses help sustain relatively stable primary productivity despite reduced irradiance, consistent with the measured NPP_14C_ during C1 (Westberry et al., 2008).

In contrast, higher light during C2 (PAR_30m_ ∼598 µmol photons m^-2^ s^-1^) resulted in photophysiological signatures of photo-stress. The maximum photochemical efficiency of PSII (Fv/Fm) decreased significantly (0.19), while Ek increased to ∼412 µmol photons m^-2^ s^-1^, consistent with reduced photosynthetic efficiency at high light and or potential iron limitation (Behrenfeld and Milligan, 2013). The antenna size decreased significantly (σ = 5), suggesting contraction of light-harvesting structures to avoid overexcitation (Schuback et al., 2017).

Elevated non-photochemical quenching (NSV = 4.52) reflects enhanced energy dissipation to protect PSII from photodamage (Lavaud et al., 2016). The faster electron turnover (1/τ) indicates increased electron cycling rates to dissipate excess excitation energy. These responses suggest that phytoplankton experienced physiological stress at higher irradiances but actively regulated energy input to maintain photochemical stability.

In C3, the continuation of C1, light levels remained moderately high (PAR_30m_ ∼559 µmol photons m^-2^ s^-1^), and the data suggest a balanced light acclimation state. Fv/Fm increased to 0.30 (+0.04 compared to C1), the highest of all cycles, and light-utilization efficiency (α) was significantly elevated. The low Ek (∼283 µmol photons m^-2^ s^-1^), together with intermediate antenna size (σ = 5.84) and low non-photochemical quenching (NSV = 2.40), suggest optimal excitation balance under these more stable light conditions. The slower electron turnover (1/τ = 0.000953 µs^-1^) reflects efficient energy conversion with minimal photoprotective dissipation. This optimized light utilization likely contributed to the relatively stable NPP_14C_ observed in C3 despite continued stratification.

During C4, light levels remained elevated (PAR_30m_ ∼579 µmol photons m^-2^ s^-1^), but photophysiological responses indicated a transitional state between high-light stress and acclimation. Fv/Fm decreased to 0.23, suggesting mild photoinhibition, while Ek remained relatively low (∼302 µmol photons m^-2^ s^-1^), indicating incomplete acclimation to higher light. The antenna size (σ = 6.17), suggests a partial enhancement of light-harvesting complexes, possibly reflecting improved nutrient availability or community shifts. Moderate non-photochemical quenching (NSV = 3.36) and rapid electron turnover rates (1/τ = 0.001397 µs^-1^) indicate active photoprotective regulation and energy balancing to maintain efficient photosynthetic operation under fluctuating light and nutrient regimes.

Overall, these photophysiological patterns illustrated how the phytoplankton community employed flexible photoacclimation strategies to maintain productivity across a range of light conditions, despite being nutrient limited (Falkowski and Raven, 2007; Suggett et al., 2009). Under low light (C1), cells maximized light capture, while under high light (C2), cells minimized excess energy input and enhanced energy dissipation. The optimal balance achieved in C3 reflects a return to more stable stratified conditions, allowing efficient light utilization without the need for strong photoprotective responses. The transitional dynamics of C4 further highlight the community’s capacity to fine-tune light harvesting and photoprotection in response to environmental shifts. These dynamic adjustments likely explain, in part, why overall primary productivity remained relatively stable across cycles, despite strong variability in surface irradiance. Flexible photoacclimation capacity is a key adaptation for phytoplankton communities inhabiting strongly stratified, oligotrophic systems such as the Argo Basin, where irradiance regimes can shift rapidly on sub-daily to multi-day timescales.

## 5. Conclusions

Across four Lagrangian experiments in the oligotrophic Argo Basin, NPP from ^14^C and N_2_-fixation in-situ incubation profiles and NPP satellite products, FRRf-derived GPP, and O_2_/Ar-based NCP measurements offer a coherent view of the productivity regime, its variability and underlying metabolic processes. Satellite and ^14^C measurements gave similar mean estimates (460 mg C m^-2^ d^-1^) of NPP. A 1.7 GPP:NPP ratio for the upper 30 m was consistent with metabolic costs of growth under low-nutrient, high-light conditions. Below 50 m, light availability, rather than nutrients, was the primary limiting factor. N_2_ fixation contributed ∼16% of NPP_14C_, accounting for the new N required to balance high export rates in the system. The general agreement among approaches supports the robustness of our rate estimates, while also highlighting the strengths and caveats of different measurements. As climate warming continues to alter surface stratification, wind forcing and nutrient supply, these detailed measurements provide a baseline for assessing future changes in N utilization and productivity in this globally important tuna spawning region.

## Acknowledgements

We thank the captain and crew of the R/V *Roger Revelle* for their outstanding support and general assistance. Research support was provided by the National Science Foundation Grants OCE-2332036 (to S.A.K.), OCE-1851381 (to K.E.S.), OCE-1851347 and OCE-1851558 (to M.R.L.), Seawater and plankton samples were collected under Australian Government permit AU-COM2021-520 and Australian Marine Parks permit PA2021-00062-2 issued by the Director of National Parks, Australia. The views expressed in this publication do not necessarily represent those of the Director of National Parks or the Australian Government.

## Author Statement

S.A.K. planned and conducted all experiments and analyzed the data. J. R. was responsible for conducting the N_2_ fixation and FRRf experiments. M.R.S. co-planned the cruise and was responsible for the in-situ array and satellite drifters, K.E.S. analyzed chlorophyll and flow cytometric samples, N.Y. contributed to deck incubations and in-situ array incubations, M.R.L. planned the cruise and permits and was responsible for nutrient sampling. S.A.K. wrote the manuscript, and all authors provided feedback on concepts in the manuscript, and provided edits of the manuscript. Generative AI was used for data-organization, coding support for Figure creation and limited text clarification revision.

## Supplemental Material

**Table S1.**
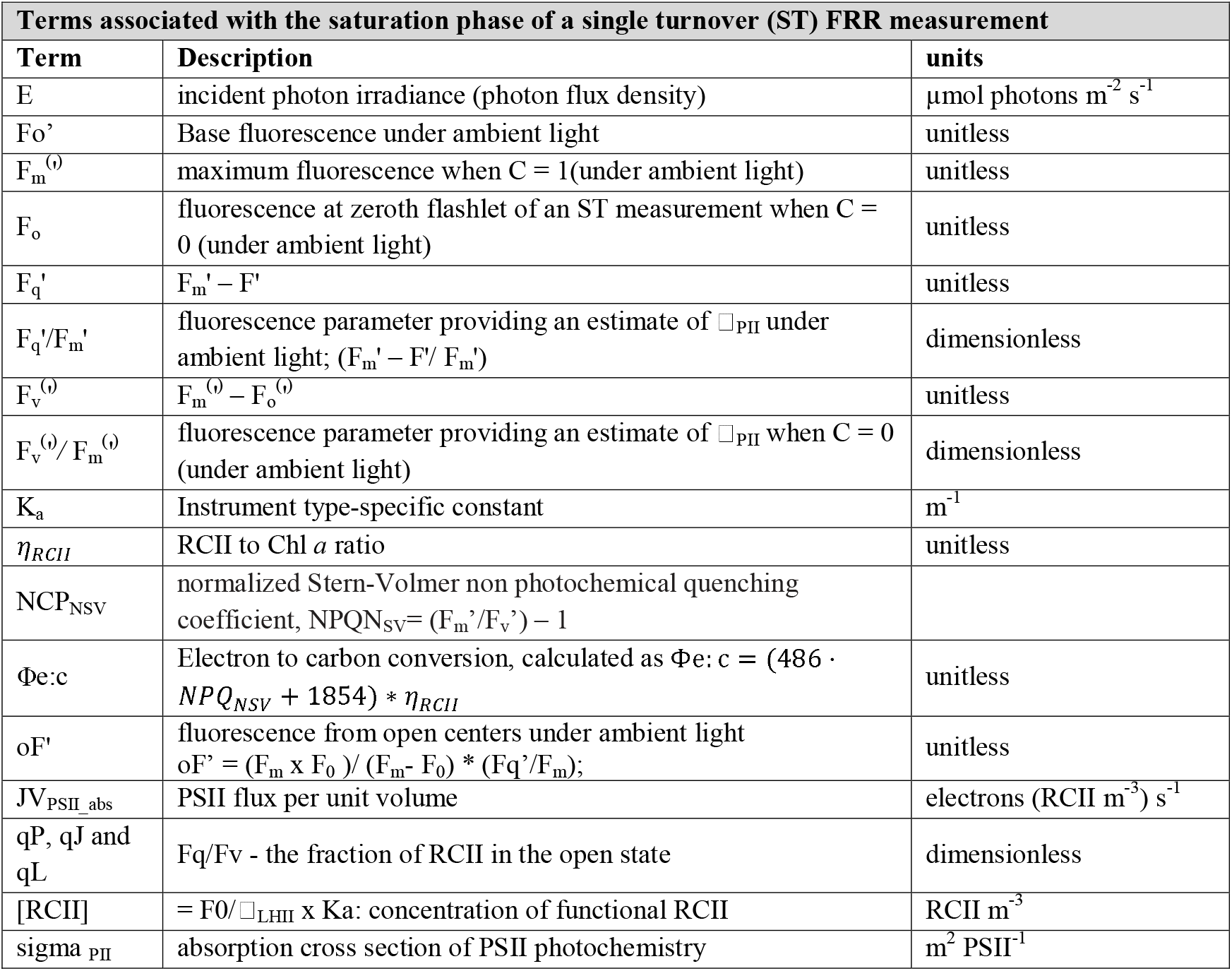
Description of terms used and units in the FRRf measurements.

**Table S2:**
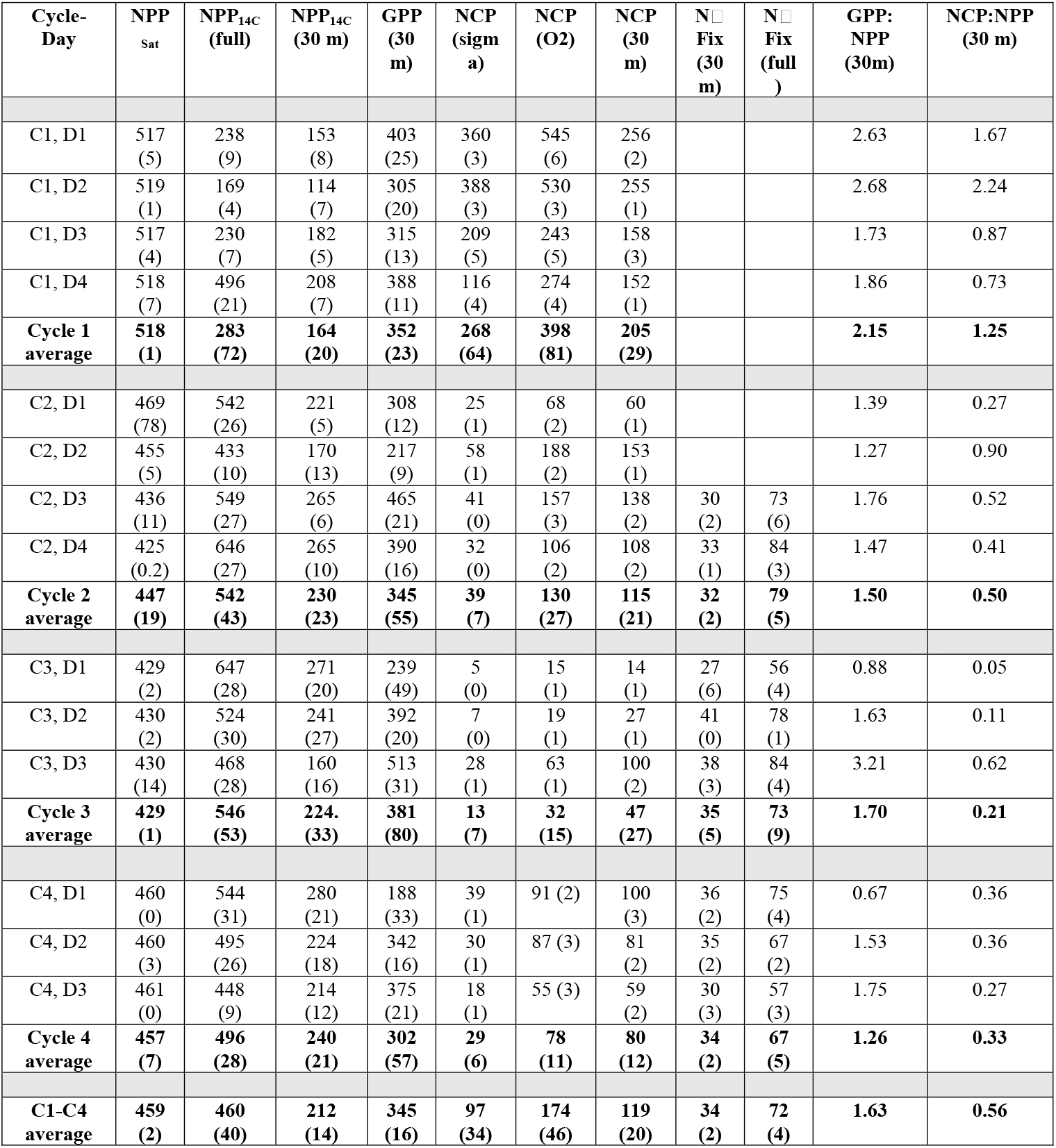
Rates of productivity measurements and ratios. Cycle (C) days (D), cycle averages and cruise averages are displayed. Rates are expressed in mg C m^-2^ d^-1^. Numbers in parenthesis denote 1 standard error of the mean.

**Table S3:** Photosynthetic parameters and light availability for each cycle. A) daily averaged surface photosynthetically active radiation integrated over the upper 30□;m (µmol photons m^-2^ s^-1^), effective quantum yield of PSII (Fv/Fm, unitless), light saturation parameter (Ek, µmol photons m^-2^ s^-1^), light utilization efficiency (α, µmol C in µmol photons m^-2^ s^-1^), functional absorption

| Cycle | PAR<br>(30 m) | Fv/Fm | Ek | $\alpha$ | $\sigma$ | NSV | $\tau$ (μs) | $1/\tau$ (ms<sup>-1</sup>) |
| --- | --- | --- | --- | --- | --- | --- | --- | --- |
| Cycle 1 | 452<br>(7.6) | 0.26<br>(0.002) | 334<br>(9.1) | 0.034<br>(0.0006) | 6.62<br>(0.07) | 2.89<br>(0.01) | 917<br>(22) | 1.25<br>(0.04) |
| Cycle 2 | 598<br>(8.8) | 0.19<br>(0.003) | 412<br>(NA) | 0.024<br>(0.0006) | 5.08<br>(0.11) | 4.52<br>(0.10) | 855<br>(20) | 1.40<br>(0.05) |
| Cycle 3 | 559<br>(10) | 0.30<br>(0.002) | 283<br>(9.8) | 0.046<br>(0.001) | 5.84<br>(0.04) | 2.40<br>(0.03) | 1120<br>(20) | 0.95<br>(0.03) |
| Cycle 4 | 579<br>(10) | 0.23<br>(0.003) | 302<br>(14) | 0.031<br>(0.0007) | 6.17<br>(0.08) | 3.36<br>(0.06) | 846<br>(20) | 1.40<br>(0.04) |

## Supplemental Figures

**Figure S1:**
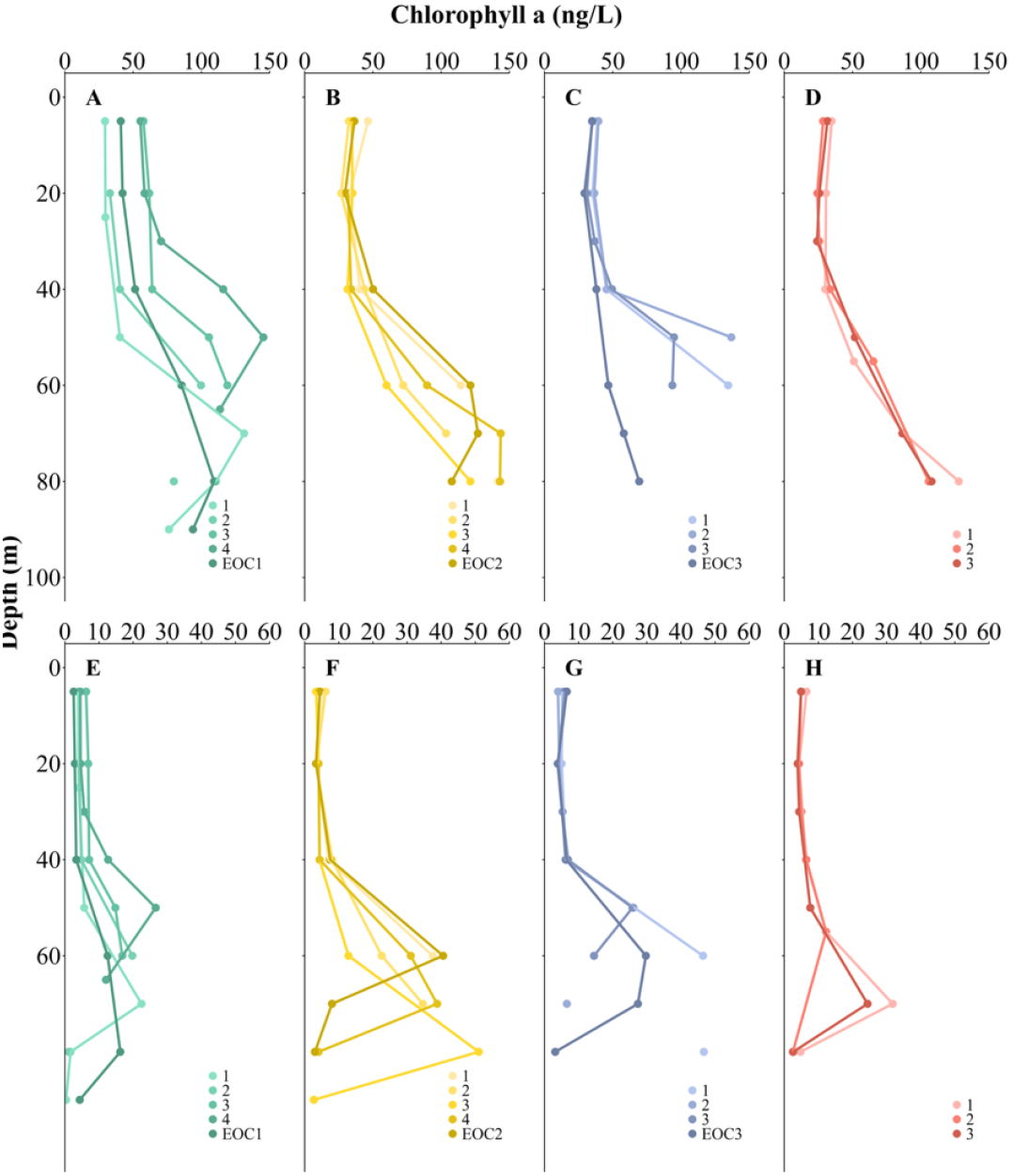
Cellular Chl*a* profiles (ng L^-1^) from each cycle from high performance liquid chromatography (HPLC), with flow cytometry used to assign monovinyl chlorophyll a to *Synechococcus* (Selph et al.).(Selph et al., this issue). A-D) C1-C4 pigment profiles (divinyl chlorophyll *a*) for *Prochlorochoccus* and E-H) C1-C4 pigment profiles (monvinyl chlorophyll *a*) for *Synechococcus*.

**Fig. S2:**
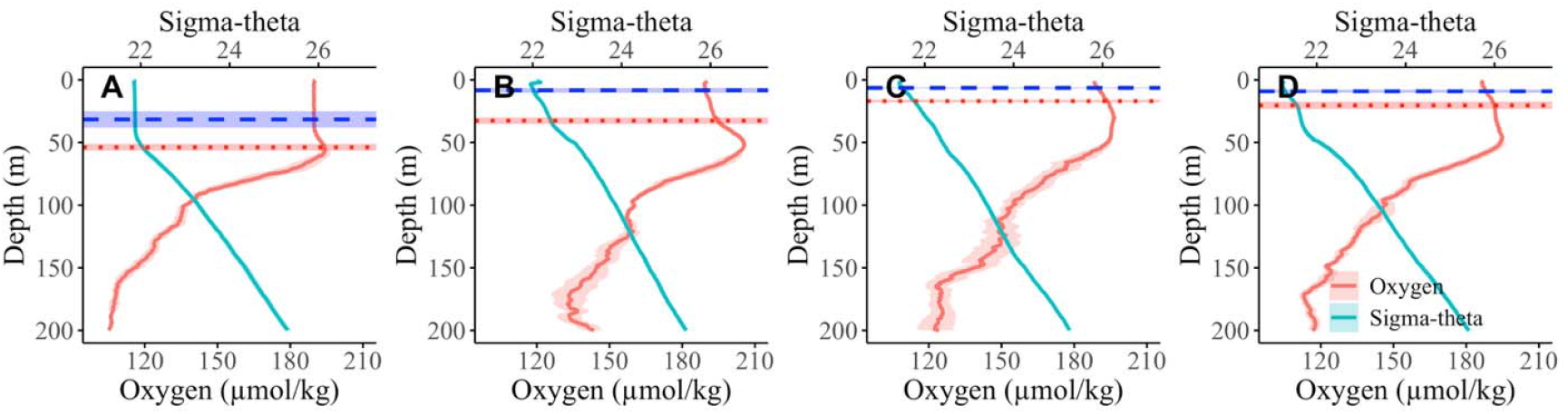
Density and O_2_ profiles for the 4 cycles. A) C1, B) C2, C) C3, D) C4. MLDs are indicated as vertical lines calculated for both Sigma-theta (Blue) and O_2_ profiles (red).

**Figure S3:**
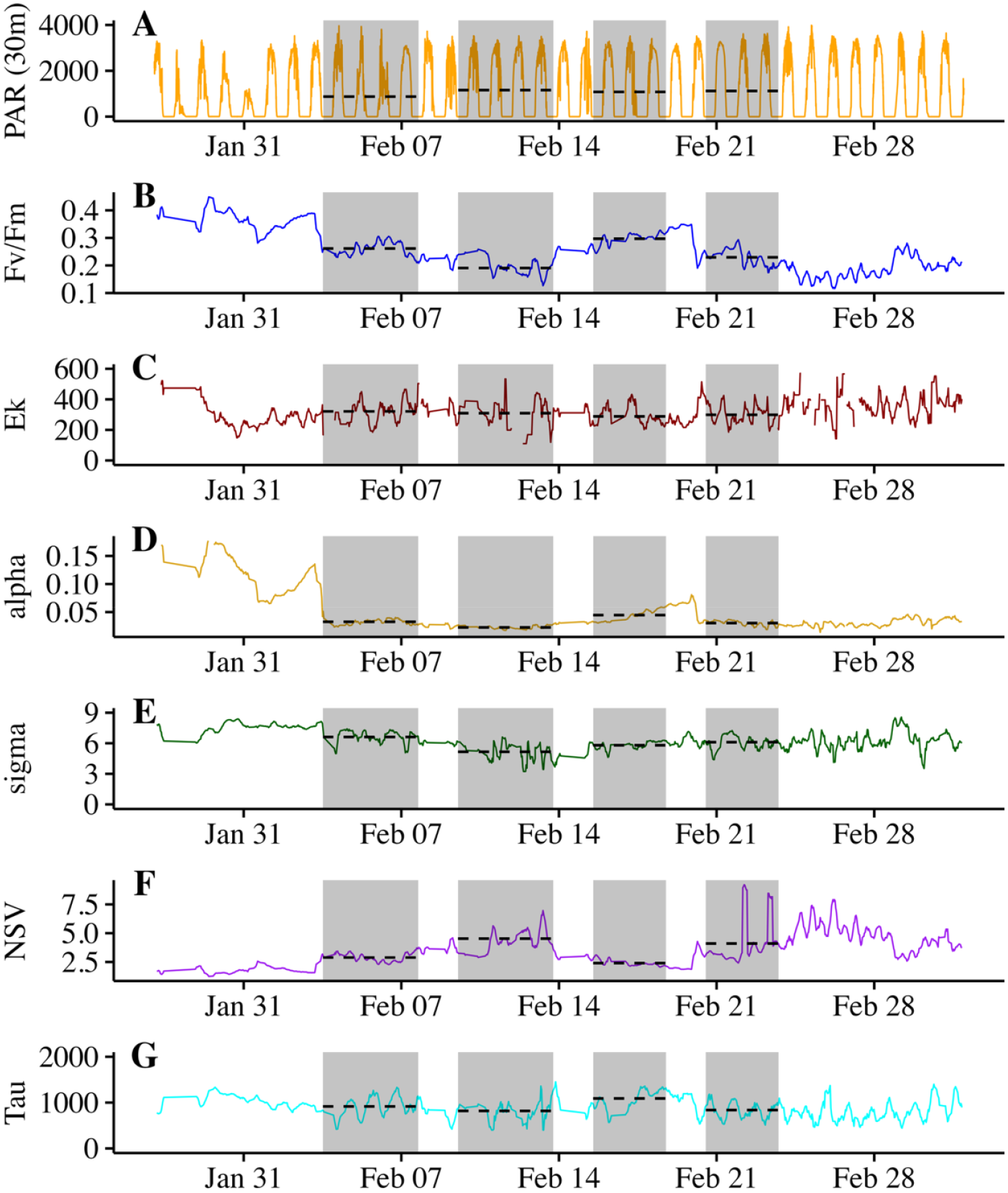
Photophysiological parameters along the BLOOFINZ cruise track. Cycles are highlighted in grey. A) Photosynthetically active radiation (PAR, µmol photons m^-2^ s^-1^); B) Photosynthetic quantum yield (Fv/Fm); C) light saturation light index (µmol photons m^-2^ s^-1^); light-limited photosynthetic efficiency (alpha), effective absorption cross-section of Photosystem II (PSII) (sigma; nm); Non photochemical quenching (expressed as normalized Stern–Volmer quenching coefficient, NSV), electron transport effective absorption cross-section of Photosystem II (PSII); turnover time of electrons through Photosystem II (Tau, µs). The horizontal lines within the cycles indicate the average values throughout the cycle.

